# Poly(ADP-ribose) potentiates ZAP antiviral activity

**DOI:** 10.1101/2020.12.17.423219

**Authors:** Guangai Xue, Klaudia Braczyk, Daniel Gonçalves-Carneiro, Daria M. Dawidziak, Katarzyna Zawada, Heley Ong, Yueping Wan, Kaneil K. Zadrozny, Barbie K. Ganser-Pornillos, Paul D. Bieniasz, Owen Pornillos

## Abstract

Zinc-finger antiviral protein (ZAP), also known as poly(ADP-ribose) polymerase 13 (PARP13), is an antiviral factor that selectively targets viral RNA for degradation. ZAP is active against both DNA and RNA viruses, including important human pathogens such as hepatitis B virus and type 1 human immunodeficiency virus (HIV-1). ZAP selectively binds CpG dinucleotides through its N-terminal RNA-binding domain, which consists of four zinc fingers. ZAP also contains a central region that consists of a fifth zinc finger and two WWE domains. Through structural and biochemical studies, we found that the fifth zinc finger and tandem WWEs of ZAP combine into a single integrated domain that binds to poly(ADP-ribose) (PAR), a cellular polynucleotide. PAR binding is mediated by the second WWE module of ZAP and likely involves specific recognition of *iso*(ADP-ribose), a repeating structural unit of PAR. Mutation of the putative *iso*(ADP-ribose) binding site in ZAP abrogates the interaction *in vitro* and diminishes ZAP activity against a CpG-rich HIV-1 reporter virus. In cells, PAR facilitates formation of non-membranous sub-cellular compartments such as DNA repair foci, spindle poles and cytosolic RNA stress granules. Our results suggest that ZAP-mediated viral mRNA degradation is facilitated by PAR, and provides a biophysical rationale for the reported association of ZAP with RNA stress granules.

## Introduction

Cells encode a variety of nucleic acid sensors that detect the presence of viral RNA or DNA by virtue of non self features or inappropriate localization. The zinc-finger antiviral protein ZAP (also known as poly(ADP-ribose) polymerase 13 or PARP13) is one such sensor and selectively binds to viral messenger RNA or viral RNA genomes [1, 2]. The ZAP-bound RNA molecules are subjected to degradation, which consequently decreases production of viral proteins and suppresses virus replication. Depending on the virus, the action of ZAP can selectively suppress viral protein expression by up to 30-fold, while cellular protein expression levels remain largely unaffected [1].

ZAP has a modular organization and is expressed as two major isoforms called ZAP-L and ZAP-S, that arise from alternative splicing and are distinguished by the presence of a C-terminal PARP (poly(ADP-ribose) polymerase)-like domain (**Fig. 1a**). Both isoforms contain an N-terminal RNA-binding domain (RBD) with four zinc fingers (here termed Z1 to Z4) that bind to CpG dinucleotides in RNA [3, 4]. Vertebrate genomes are depleted of CpG content, and it is the relative scarcity of this dinucleotide in cellular RNA compared to susceptible viral RNA that explains selective ZAP-mediated degradation [5]. A truncated ZAP fragment (here called ZAP-N; **Fig. 1a**) that essentially consists of only the RBD is both necessary and sufficient for directing viral RNA degradation [1]. However, there are indications that other ZAP domains are also important for its antiviral function [6, 7].

**Fig. 1.**
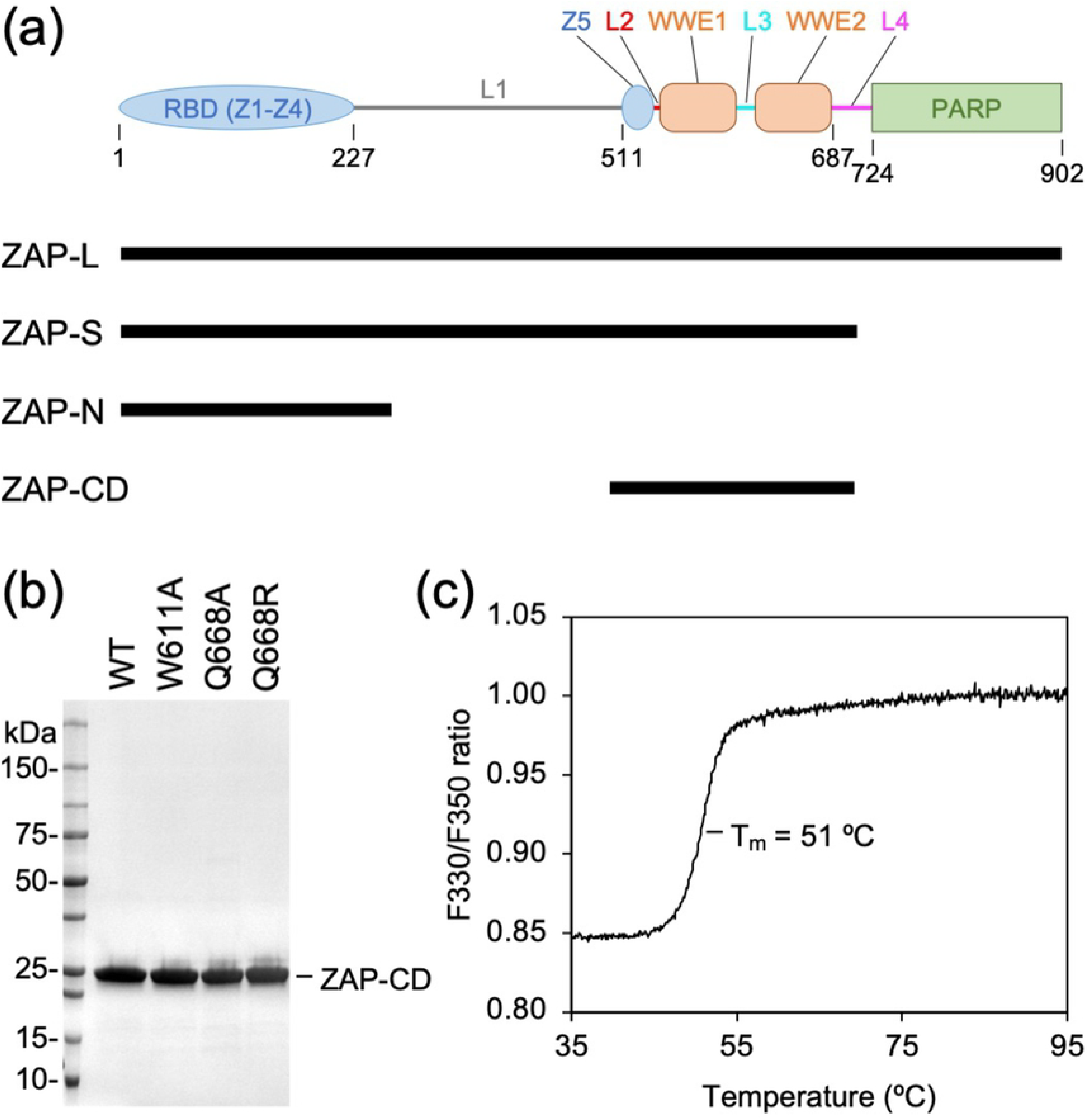
Modular domain organization of ZAP. (**a**) Domain diagram of the ZAP primary sequence. Modules are colored according to their structural properties (zinc fingers Z1, Z2, Z3, Z4 and Z5 in blue; WWE1 and WWE2 domains in orange; PARP in green; inter-domain linkers L1, L2, L3, and L4 in gray, red, cyan and magenta). Indicated below are the two major naturally-occurring splice isoforms (ZAP-L and ZAP-S), the minimally active antiviral fragment (ZAP-N) [1], and the central domain described in this study (ZAP-CD). (**b**) SDS-PAGE profiles of purified recombinant ZAP-CD proteins used in this study. (**c**) Differential scanning fluorimetry profile of wild type (WT) ZAP-CD shows a single transition. The apparent melting temperature (*T*_m_) is 50.9 ± 0.1 °C (determined with five independent protein preparations).

In both ZAP-L and ZAP-S, the RBD is connected by a long linker segment to a fifth zinc finger (Z5) and two WWE domains (WWE1 and WWE2) (**Fig. 1a**). These additional ZAP domains have unknown function, but WWE domains in other proteins are reported to have a general role in binding to poly (ADP-ribose) (PAR) [8, 9]. PAR is a cellular polynucleotide that has been shown to function as a scaffold or collective docking site for multiple protein partners, thereby allowing for sustained co-localization of the components of cellular pathways [10, 11]. Here, we show that Z5, WWE1 and WWE2 are sub-domains or modules that integrate into a composite fold, which we term the ZAP central domain (ZAP-CD). Structural and biochemical analyses revealed that ZAP-CD binds to PAR through the second WWE module. Both ZAP [12–15] and PAR [16, 17] have been previously reported to localize to so-called RNA stress granules, which constitute a type of non-membranous cytoplasmic compartment that facilitates RNA turnover and antiviral responses [18, 19]. Our studies suggest that PAR may coordinate the stable co-localization of ZAP and its co-factors in stress granules and thereby facilitate efficient recognition and/or degradation of ZAP-bound RNA.

## Results

### ZAP Z5, WWE1 and WWE2 form a single composite domain

We first aimed to define a protein construct from the central regions of ZAP that could be characterized biochemically. The Z5, WWE1 and WWE2 modules were insoluble when individually overexpressed recombinantly in *E. coli*, but a ZAP construct spanning residues 498-699 and containing all three was highly soluble and could be purified to homogeneity (**Fig. 1b**). The purified ZAP-CD protein exhibited a single folding transition in thermal melting experiments (**Fig. 1c**). We then determined the crystal structure of ZAP-CD to 2.5 Å resolution (*R*_work_/*R*_free_ = 0.22/0.26) (**Fig. 2a** and **Table 1**). The Z5, WWE1 and WWE2 modules each form a compact fold or sub-domain. Close packing between the three modules is mediated by well-ordered “linker” residues, which we term L2, L3 and L4 (colored in red, cyan and magenta in **Fig. 2a**); although separated in sequence these linkers come together in the middle of the structure and glue together the three sub-domains. Thus, the three modules or sub-domains of ZAP-CD are integrated into a composite fold that likely behaves as a single functional unit.

**Fig. 2.**
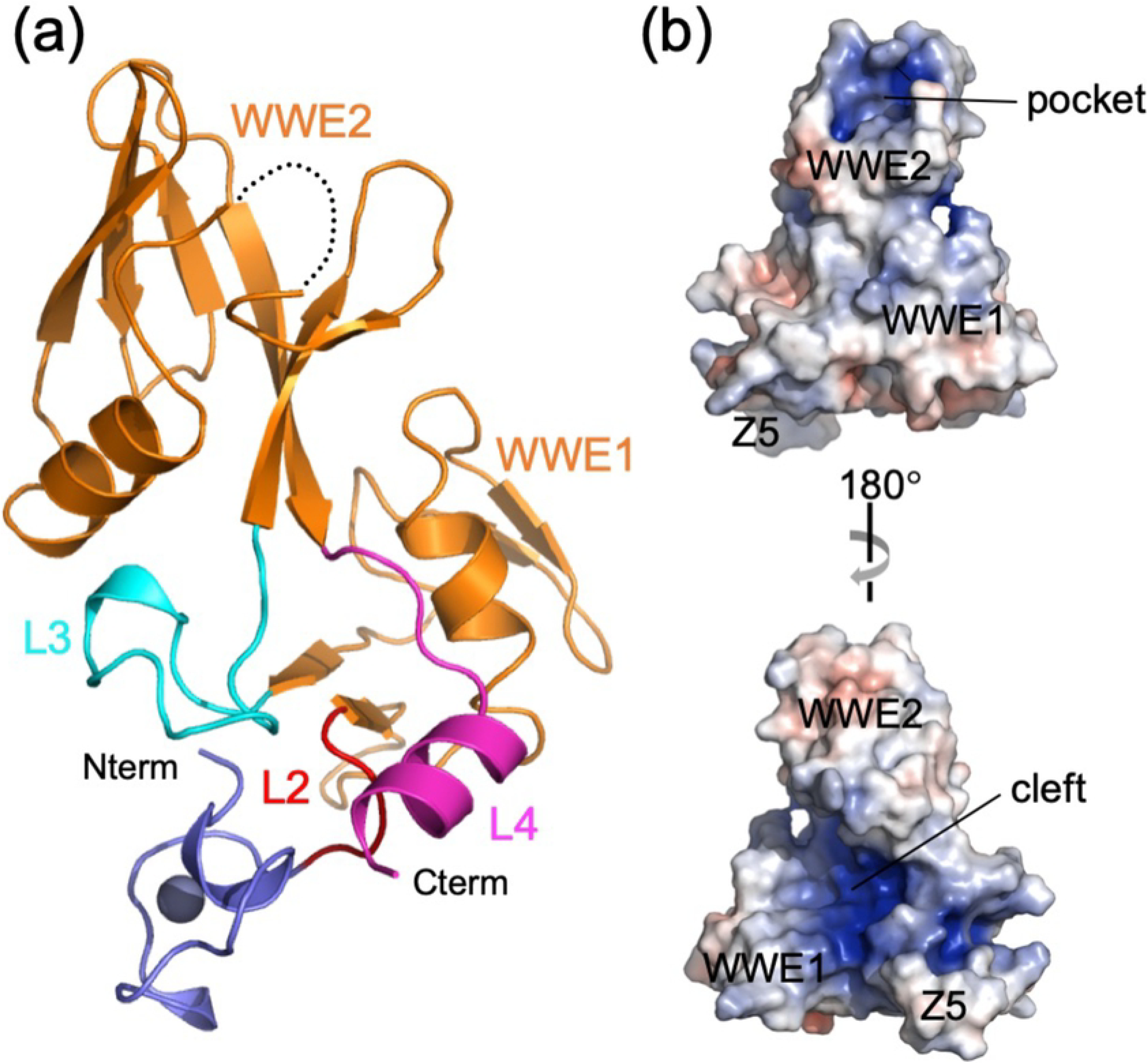
Crystal structure of ZAP-CD. (**a**) Ribbons representation. Modules are colored according to **Figure 1a** (Z5 in blue, WWE1 and WWE2 in orange), as are the linkers (L2 in red, L3 in cyan, and L4 in magenta). The amino and carboxyl termini are also indicated. Dashed black lines denote a disordered loop in the second WWE module. (**b**) Orthogonal surface views colored according to electrostatic potential from red (negative) to blue (positive). The Z5, WWE1 and WWE2 modules are indicated, as are the putative *iso*(ADP-ribose) pocket (top) and the extended electropositive cleft (bottom).

**Table 1.** Crystallographic statistics.

| SeMet ZAP-CD |  |
| --- | --- |
| **Diffraction Data** |  |
| Beamline | APS 22ID |
| File root | ID22_032818_pck4pn2_SeMetZAP11 |
| Wavelength (Å) | 0.97856 |
| Processing program | HKL2000 |
| Space group | P321 |
| Cell dimensions | $a = b = 89.637 \text{ Å}$, $c = 53.055 \text{ Å}$<br>$\alpha = \beta = 90^\circ$, $\gamma = 120^\circ$ |
| Resolution range, Å | 50-2.50 (2.54-2.50) |
| $R_{\text{sym}} / R_{\text{meas}} / R_{\text{pim}}$ | 0.14 (1) / 0.18 (1) / 0.03 (0.24) |
| CC1/2 | 0.998 (0.893) |
| Mean $I/\sigma(I)$ | 24.2 (5.0) |
| Completeness, % | 99.5 (100) |
| Average redundancy | 48.0 (24.1) |
| Wilson B-factor, Å<sup>2</sup> | 35.6 |
| **SAD Phasing** |  |
| Phasing program | PHENIX (phenix.autosol) |
| No. of Se sites | 5 |
| Figure of merit | 0.38 |
| **Refinement Statistics** |  |
| Refinement program | PHENIX (phenix.refine) |
| Resolution range | 39-2.50 (2.70-2.50) |
| No. of unique reflections | 8,296 (1,701) |
| Reflections in free set | 428 (89) |
| $R_{\text{work}} / R_{\text{free}}$ | 0.22 (0.26) / 0.26 (0.29) |
| No. of nonhydrogen atoms |  |
| protein | 1,447 |
| zinc | 1 |
| water | 33 |
| Average B-factor (Å<sup>2</sup>) |  |
| protein | 42.1 |
| zinc | 43.2 |
| water | 34.7 |
| Coordinate deviations |  |
| bond lengths, Å | 0.003 |
| bond angles, ° | 0.530 |
| **Validation and Deposition** |  |
| Ramachandran plot |  |
| favored, % | 97.8 |
| outliers, % | 0 |
| MolProbity clashscore | 0.71 |
| PDB ID | 7KZH |

Values in parenthesis are for the highest resolution shell.

Examination of the surface features and electrostatic potential of ZAP-CD revealed two major regions of interest: a deep pocket in the second WWE module (**Fig. 2b**, top), which appeared suitable for binding an aromatic ligand (discussed in more detail below), and a deep cleft or ridge running along one side of the composite domain (**Fig. 2b**, bottom). A similar cleft was observed in the tandem WWE fold of Deltex, which was proposed to bind to extended polypeptide segments [20]. The highly electropositive nature of the cleft suggests that it may also be well suited to bind negatively-charged, non-proteinaceous polymers such as polynucleotides.

### ZAP-CD contains a putative high-affinity PAR binding site

The WWE domain was first described as an independent protein-folding module with a characteristic signature of conserved tryptophan, glutamate and arginine residues, and is found in many proteins that function in ubiquitination and PARylation pathways [8, 9]. In ZAP, the two WWE modules share the same canonical β-strand/α-helix fold as expected, but differ considerably with regards to the loops connecting the β-strands (compare **Fig. 3a** and **Fig. 3b**). In the second WWE module, the loops are more extended and generate the abovementioned pocket (**Fig. 3b**). The walls of the pocket are lined by hydrophobic sidechains (W611, Y621, Y659), and at the bottom of the pocket is a partially buried glutamine residue (Q668). Comparison of ZAP’s second WWE module to that of the single WWE in RNF146 (an E3 ubiquitin ligase involved in DNA repair) [21] (**Fig. 3c**) revealed that these pocket sidechains are highly conserved in both identity and three-dimensional configuration, suggesting a common function (**Fig. 3d**). In RNF146, the pocket is a nM-affinity binding site for *iso* (ADP-ribose), a repeating structural unit of PAR. In the RNF146 crystal structure with bound *iso* (ADP-ribose), the buried glutamine (Q153, equivalent to Q668 in ZAP) makes hydrogen bonds with the adenine ring of the ligand; a hydrophobic sidechain (Y107, equivalent to W611 in ZAP) makes a *pi*-stacking interaction with the same adenine ring [21]. Residues that mediate binding to the phosphate groups are likewise conserved or highly similar between ZAP and RNF146 (e.g., R163 in RNF146 and K677 in ZAP) (**Fig. 3e**). In the case of ZAP-CD, the protein construct was crystallized without added ligand, but closer examination of the electron density maps within the pocket revealed well-defined residual difference densities (green and blue mesh in **Fig. 3b**), whose shapes were consistent with a bound adenine ring and a phosphate. Indeed, protein backbone-guided superposition of the ZAP-CD and RNF146 structures places the RNF146-bound *iso* (ADP-ribose) ligand precisely within these densities (**Fig. 3e**). This close correspondence indicates that, like RNF146, ZAP is a PAR-binding protein. Note that *iso* (ADP-ribose) does not naturally exist in cells and was originally synthesized as a reagent to establish the specificity determinant of RNF146 for PAR [21]. We surmise that some abundant small molecule containing an adenine ring and phosphate groups (e.g., ATP, ADP-ribose or similar) co-purified and co-crystallized with our recombinant ZAP-CD protein (see also Materials and Methods). In any case, the important point is that the above analyses strongly indicated that, just like the RNF146 WWE domain, the second WWE module of ZAP binds PAR.

**Fig. 3.**
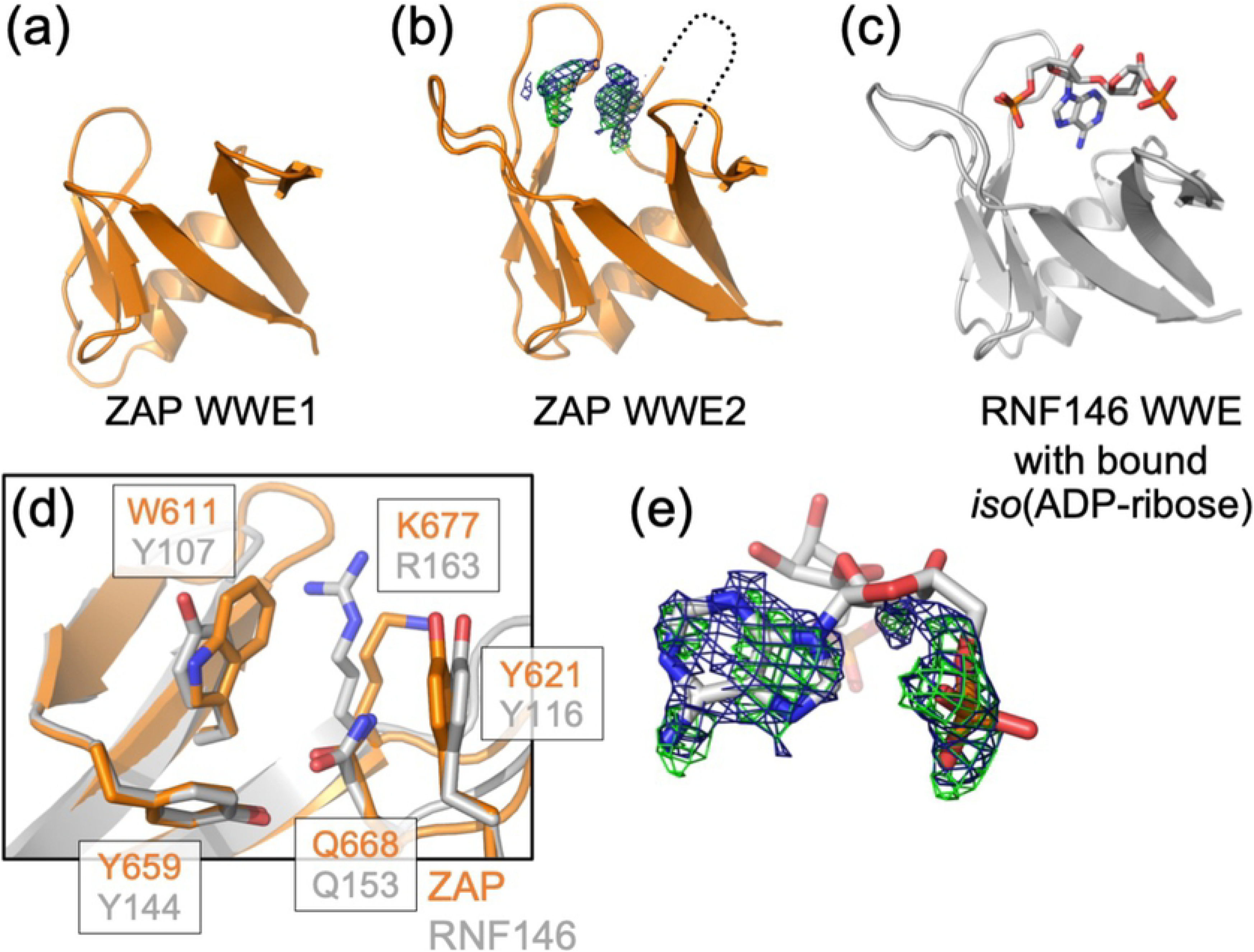
Structural analysis of the WWE modules. (**a-c**) Comparison of ZAP WWE1 (a), ZAP WWE2 (b) and RNF146 WWE with bound *iso*(ADP-ribose) ligand (PDB 4QPL) [21] (c). The three structures are shown in the same orientation. (**d**) Backbone-guided superposition of the *iso*(ADP-ribose) binding pockets in ZAP-CD (orange) and RNF146 (gray). Equivalent sidechains surrounding the pocket are shown explicitly and labeled. (**e**) Same superposition as (d) but showing only the residual difference densities observed in the ZAP-CD structure and the iso(ADP-ribose) ligand in the RNF146 structure. Note the excellent shape and positional matches to the adenine ring and one of the two phosphate groups. Green mesh in (b) and (e) represents unbiased m*F*_o_-D*F*_c_ density contoured at 2σ; blue mesh represents 2m*F*_o_-D*F*_c_ density contoured at 1σ.

### ZAP-CD binds PAR *in vitro*

To directly test whether ZAP interacts with PAR, we enzymatically synthesized and purified PAR polymers *in vitro* [21, 22] (**Fig. 4a**). We prepared two forms of PAR, which we term ‘high-MW PAR’ and ‘low-MW PAR’ to reflect their elution behavior from a preparative sizing column (**Fig. 4b**) and electrophoretic migration in an agarose gel (**Fig. 4c**). Note that in both of these preparations the PAR polymers are heterogeneous in length and probably constitute both linear and branched forms.

**Fig. 4.**
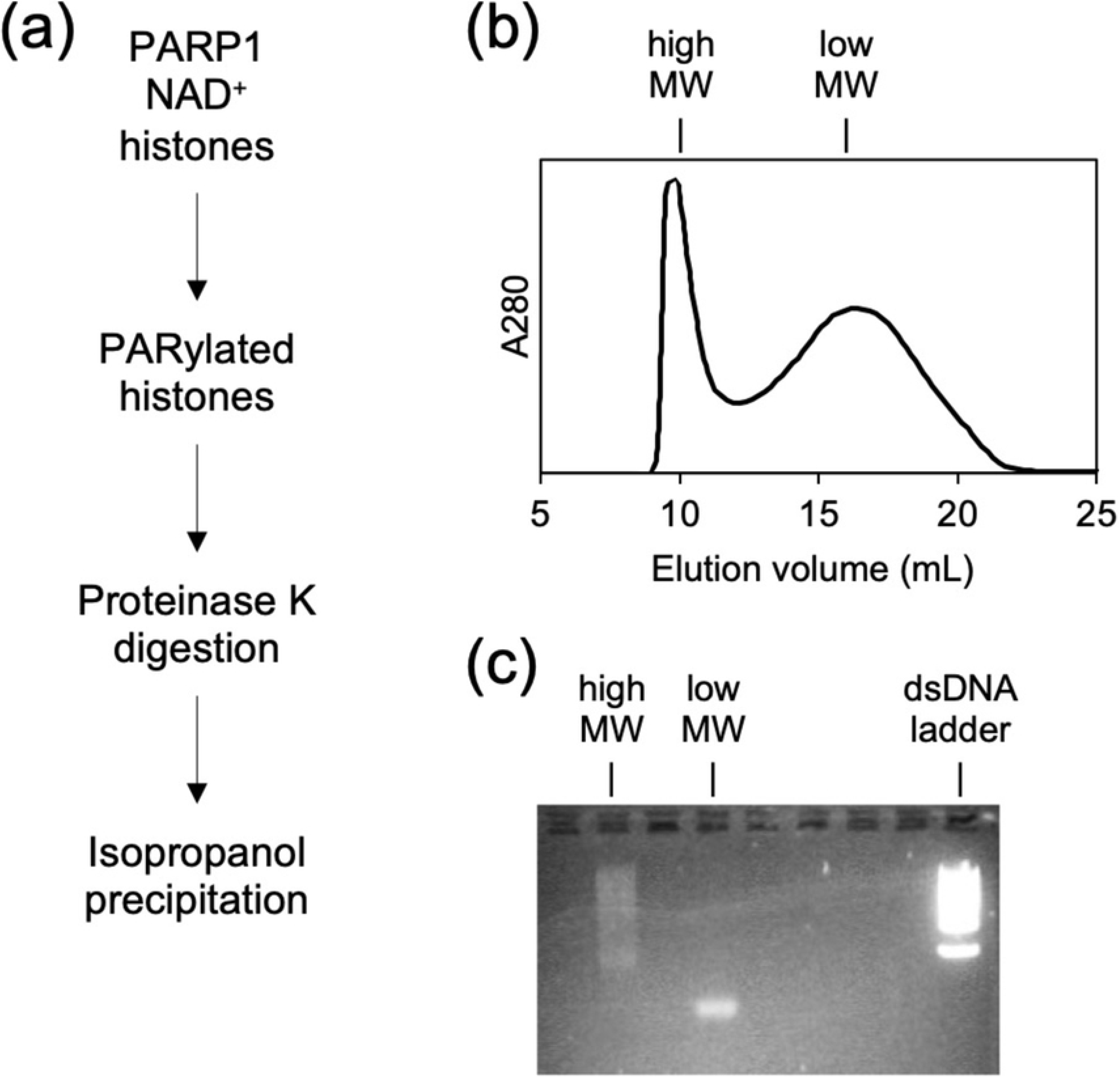
Synthesis and preparation of PAR. (**a**) Histones were PARylated by incubation with recombinant PARP1 enzyme and NAD^+^ [22]. After proteolytic digestion to remove the proteins, released PAR polymers were purified by isopropanol precipitation. (**b**) Size exclusion profile on a preparative Superdex 200 column, after resuspension of the isopropanol precipitate. (**c**) Agarose gel electrophoresis profiles of the indicated fractions. Note that even though the ‘low-MW’ fraction appears as a single band, it is still a mixture of different lengths (and likely includes branched forms) of PAR.

We then used analytical size exclusion chromatography to test for a binding interaction (**Fig. 5a**). In this experiment, a positive interaction can be generally expected to manifest in one of two ways. High-affinity binding can result in formation of a stable complex that elutes with an apparent size (strictly speaking, hydrodynamic radius) greater than the early-eluting component. Less stable complexes can dissociate and exchange with the unbound components during the chromatography run to generate an elution profile in which the late-eluting component peak is smeared towards an earlier volume. Initially, we tested for binding between ZAP-CD and low-MW PAR. In control experiments, ZAP-CD alone eluted as a single peak from an analytical Superdex 200 column with an elution volume of ~18 mL (**Fig. 5a**, green curve). Low-MW PAR eluted at an earlier volume of ~14.5 mL from the same column (**Fig. 5a**, black curve). When the two components were mixed prior to sample injection, both types of elution behavior described above were observed. The profile consisted of a peak with an elution volume of ~13.5 mL, which is earlier than either PAR or ZAP-CD alone and indicative of a stable complex; in addition, the trailing ZAP-CD peak was also smeared towards earlier elution volumes (**Fig. 5a**, orange curve). SDS-PAGE confirmed the presence of protein in all the relevant fractions (insets in **Fig. 5a**). These results show that ZAP-CD indeed binds PAR *in vitro*, and that furthermore, two types of ZAP-CD/PAR interactions can be discerned from the exchange behavior of the complexes during size exclusion chromatography.

**Fig. 5.**
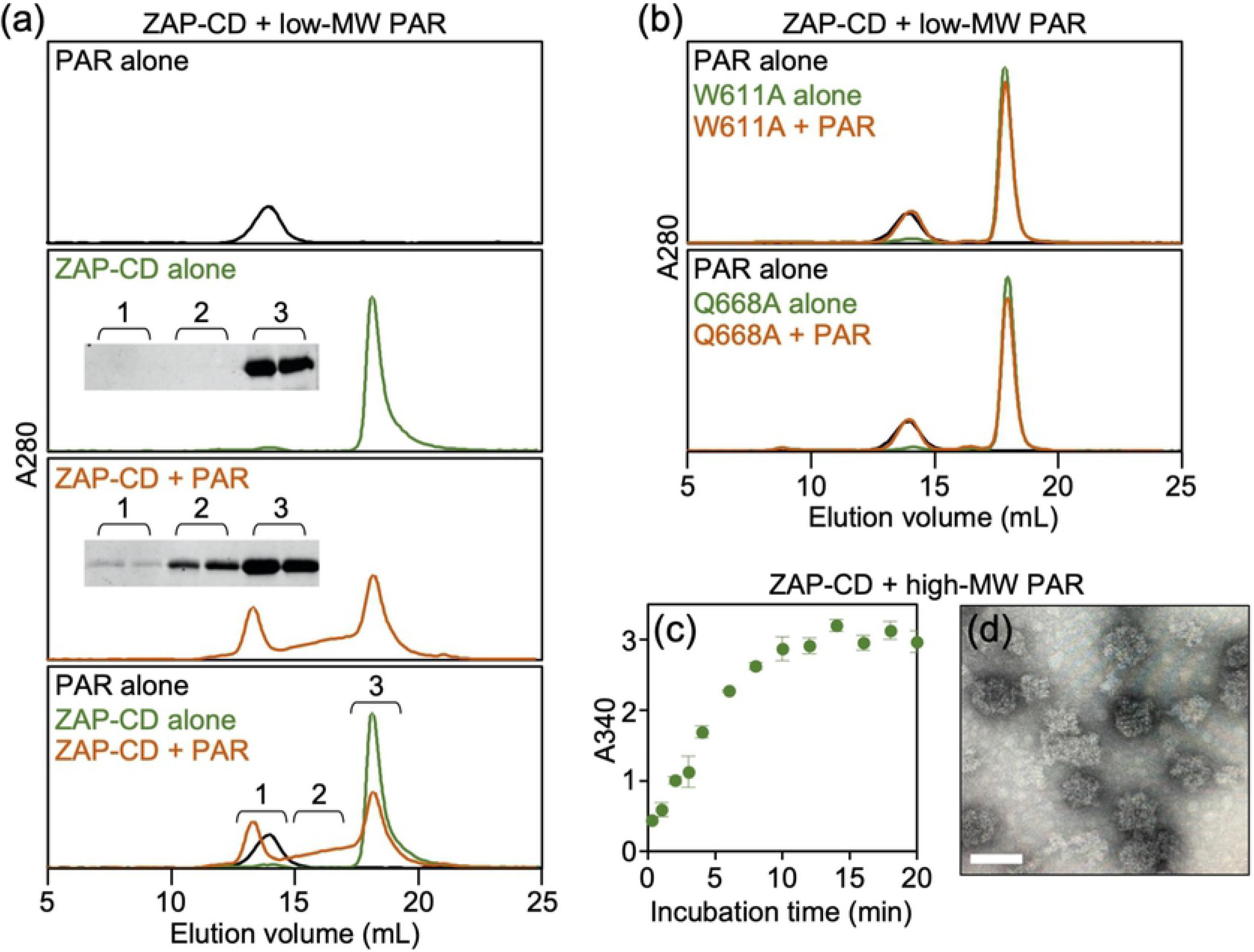
ZAP-CD binds to PAR *in vitro*. (**a**) Size exclusion binding assay with purified ZAP-CD and low-MW PAR. The three top panels show individual analytical Superdex 200 size exclusion profiles of purified PAR alone (black), ZAP-CD alone (green) and mixed ZAP-CD and PAR after 20 min incubation (orange). The bottom panel shows an overlay of all three curves. Insets show SDS-PAGE analysis of fractions indicated in the bottom panel. Results are representative of two independent experiments, each done in two replicates. (**b**) Representative results of assays performed with low-MW PAR and the indicated ZAP-CD mutants. Results are representative of two independent experiments. (**c**) Light scattering of a binding reaction containing ZAP-CD and high-MW PAR as a function of incubation time. Error bars indicate the standard deviation of three independent measurements. (**d**) Negative stain electron microscopy image of ZAP-CD/high-MW PAR complexes (20 min time point in c). Scale bar, 100 nm.

As described above, the ZAP-CD structure revealed a putative *iso* (ADP-ribose) binding pocket that contains a buried glutamine residue, Q668, surrounded by hydrophobic sidechains including W611 (**Fig. 3d**). To confirm the importance of this pocket for PAR binding, we purified and tested ZAP-CD proteins harboring W661A, Q668A or Q668R mutations for binding (**Fig. 1b**). The mutant proteins did not bind low-MW PAR as evidenced by the elution profiles of the mixed samples, which were simple sums of the profiles of the individual components (**Fig. 5b**). These results confirm that the shifts in elution volume observed in **Figure 5a** arise from specific interactions between ZAP-CD and PAR, and that these interactions involve binding of the *iso* (ADP-ribose) unit of PAR to the WWE pocket, analogous to RNF146. Furthermore, both the non-exchanging and exchanging ZAP-CD/low-MW PAR complexes require the *iso* (ADP-ribose) pocket.

We next tested binding of ZAP-CD to the high-MW PAR preparation. Mixing of protein and polynucleotide resulted in rapid solution turbidity, as evidenced by an increase in the solution light scattering signal as a function of incubation time (**Fig. 5c**). Negative stain electron microscopy revealed that the sample consisted of large globular particles around 100 nm in size (**Fig. 5d**). Given the polynucleotide nature of PAR, we expected these particles to have filamentous character, and indeed, the individual globules appear to be compacted filaments, although this requires further confirmation. Interestingly, the ZAP-CD/high-MW PAR particles are similar in size and appearance to PAR polymers and PARylated PARP enzymes observed in previous electron microscopy studies [23–25]. Our results suggest that ZAP and PAR may form higher-order complexes or assemblies in cells.

### ZAP-CD binds PAR in cells

To confirm that ZAP interacts with PAR in cells, we overexpressed HA-tagged ZAP proteins in HEK 293T cells and performed co-immunoprecipitation experiments (**Fig. 6**). HA-tagged ZAP-CD efficiently co-precipitated PAR from clarified cell lysates (**Fig. 6a**, lane 2). In contrast, ZAP-CD proteins harboring the Q668A or Q668R mutations in the putative *iso* (ADP-ribose) pocket did not co-precipitate PAR (**Fig. 6a**, lanes 3 and 4). Experiments performed with HA-tagged ZAP-L, a naturally-occurring full-length isoform of ZAP, likewise revealed that ZAP-L could co-precipitate PAR (**Fig. 6b**, lane 2). In contrast to the shorter ZAP-CD construct, however, the Q668A and Q668R ZAP-L mutants still co-precipitated PAR, although at appreciably lower amounts than wild type ZAP-L (**Fig. 6b**, compare lanes 3 and 4 with lane 2). These results indicate that the ZAP central domain indeed binds to PAR in cells as predicted by our structural and biochemical analyses, but that other domains in ZAP-L may also independently associate with PAR (see Discussion).

**Fig. 6.**
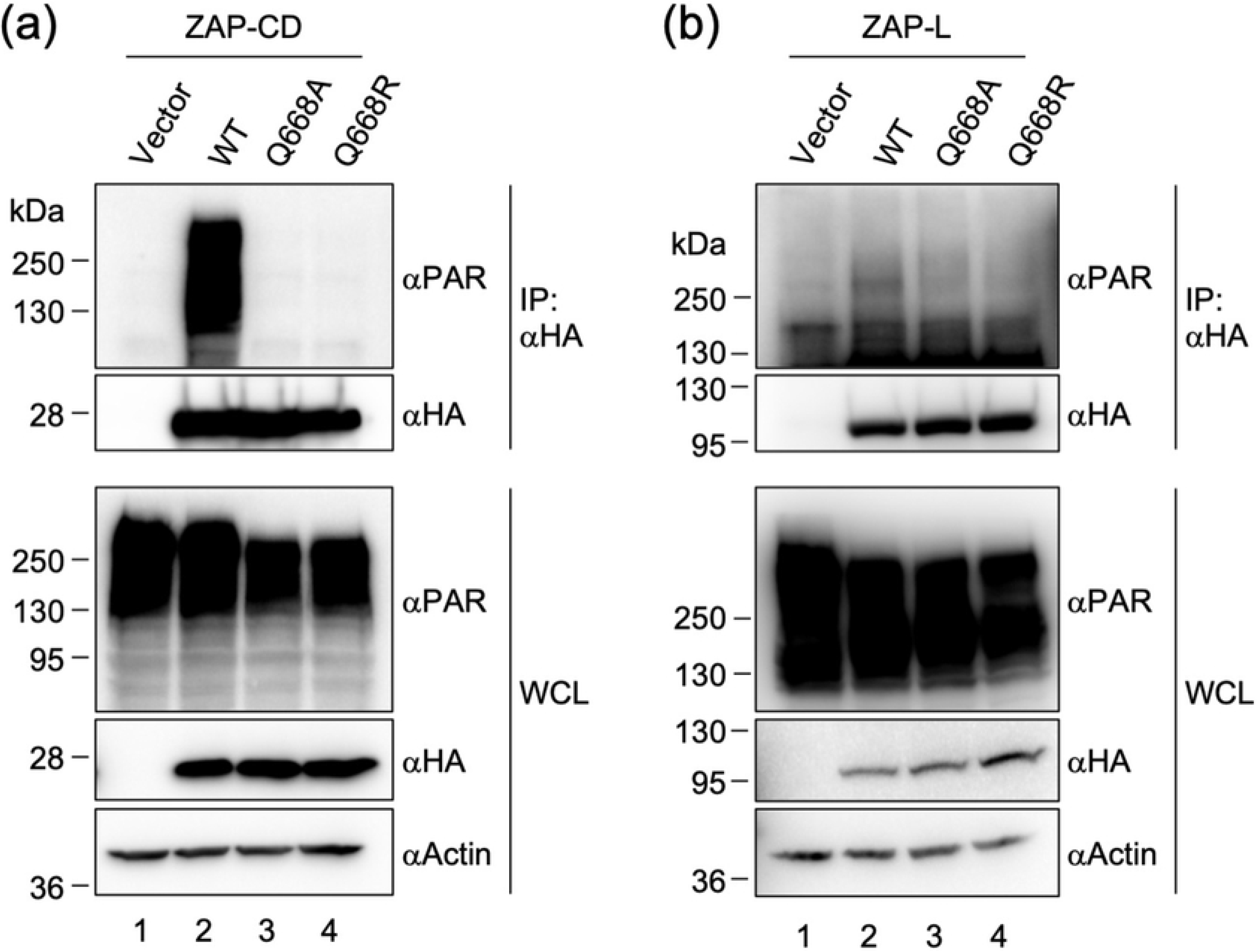
PAR co-immunoprecipitates with ZAP. (**a-b**) HEK 293T cells were transfected with empty vector or the indicated HA-tagged ZAP-CD constructs (a) or ZAP-L constructs (b). Forty-eight hours later, whole cell lysates (WCL) were subjected to pull-down with anti-HA antibody followed by immunoblotting with the indicated antibodies. Actin was used as loading control. Results are representative of three (ZAP-CD) or two (ZAP-L) independent experiments.

PAR is ubiquitously found in cells but becomes enriched within non-membranous sub-cellular compartments called RNA stress granules upon stress induction or virus challenge. For example, treatment of cells with arsenite (which causes oxidative stress) induces PAR accumulation in stress granules [16]. We took advantage of this property to directly examine association of ZAP and PAR in the cellular setting. HA-tagged ZAP-CD was expressed in HeLa cells under conditions that allowed facile visualization of both ZAP-CD and PAR by immunofluorescence microscopy (**Fig. 7**). In untreated cells, both ZAP-CD and PAR showed diffuse staining (**Fig. 7a**, left panels). Upon treatment of the cells with 0.25 mM sodium arsenite, a fraction of the PAR staining redistributed into punctate accumulations (indicated by white arrows in **Fig. 7a**, right, middle panel) that were clearly distinguished from the diffuse background fraction. (Control experiments confirmed that these puncta also contained established stress granule markers.) Arsenite treatment also induced the redistribution of ZAP-CD (**Fig. 7a**, right, top), and importantly the punctate ZAP-CD accumulations co-localized with the PAR accumulations (**Fig. 7a**, right, bottom). In contrast, the ZAP-CD Q668R mutant did not re-localize with PAR (**Fig. 7b**). We therefore conclude that the ZAP central domain binds PAR, both *in vitro* and in cells.

**Fig. 7.**
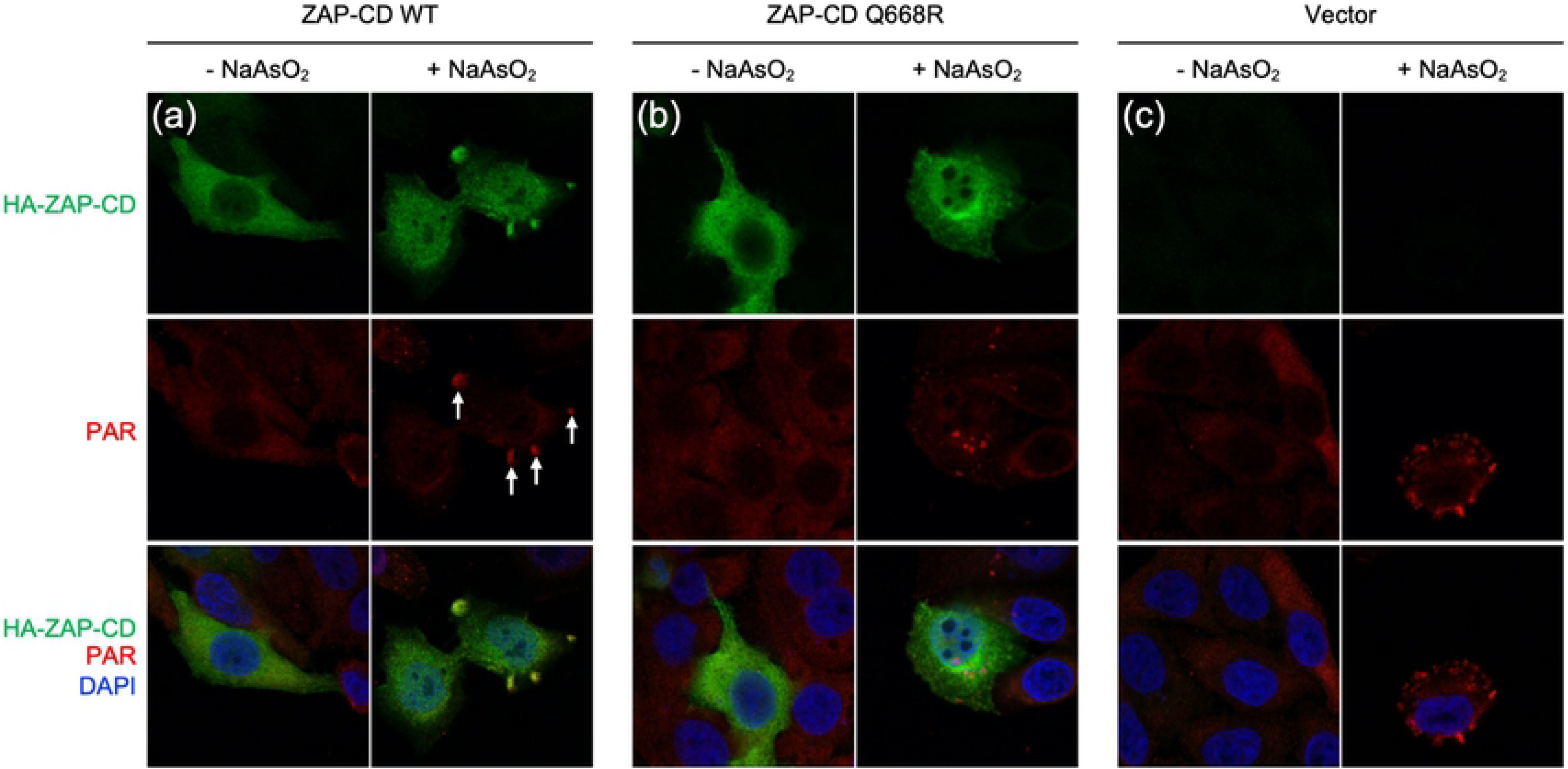
ZAP co-localizes with PAR in cytoplasmic puncta. HeLa cells were transiently transfected with (**a**) vector encoding HA-tagged ZAP-CD, (**b**) vector encoding HA-tagged ZAP-CD Q668R mutant, or (**c**) empty vector control. Twenty-four hours later, cells were treated with sodium arsenite or mock-treated, fixed, immunostained with anti-HA and anti-PAR primary antibodies followed by dye-conjugated secondary antibodies, and imaged by using fluorescence microscopy. Results are representative of two independent experiments.

### PAR-binding by the central domain potentiates ZAP antiviral activity

Having established the biochemical properties of the ZAP central domain both *in vitro* and in cells, we next tested whether its PAR binding activity would affect ZAP’s antiviral function against CpG-enriched HIV-1. We used an engineered HIV-1 mutant, termed (NL4.3 CG-High), that was previously generated by synonymous mutagenesis to contain a higher number of CpGs compared to wild type HIV-1 [5]. ZAP directly binds to and directs the degradation of the CpG-rich viral RNA transcripts, thereby reducing viral protein synthesis and the yield of (NL4.3 CG-High) virus. We transfected ZAP-deficient HEK293T cells with proviral plasmids encoding wild type HIV-1 control (NL4.3 WT) or CpG-enriched virus (NL4.3 CG-High), together with varying amounts of expression vectors encoding either wild type ZAP-L or the ZAP-L Q668R mutant. Virus yields were measured 48 h after transfection. While both the wild type and CpG-enriched viruses gave similar yields in the absence of ZAP-L, progressively increasing the expression of ZAP-L resulted in corresponding reduction in the yield of infectious units of HIV-1 (NL4.3 CG-High) but not (NL4.3 WT) (**Fig. 8a**). Notably, the PAR-binding deficient ZAP-L mutant (Q668R) exhibited a 5- to 10-fold reduced antiviral potency when compared to wild type ZAP-L. The diminished antiviral activity was also reflected in the amount of Env protein synthesized in transfected cells (gp160 and gp120, **Fig. 8b**). These results confirm that the PAR-binding property of the central domain is not essential for ZAP-L-mediated inhibition of CpG-enriched HIV-1 replication. Nevertheless, loss of the ability to bind PAR correlated with an appreciable decrease in antiviral activity.

**Fig. 8.**
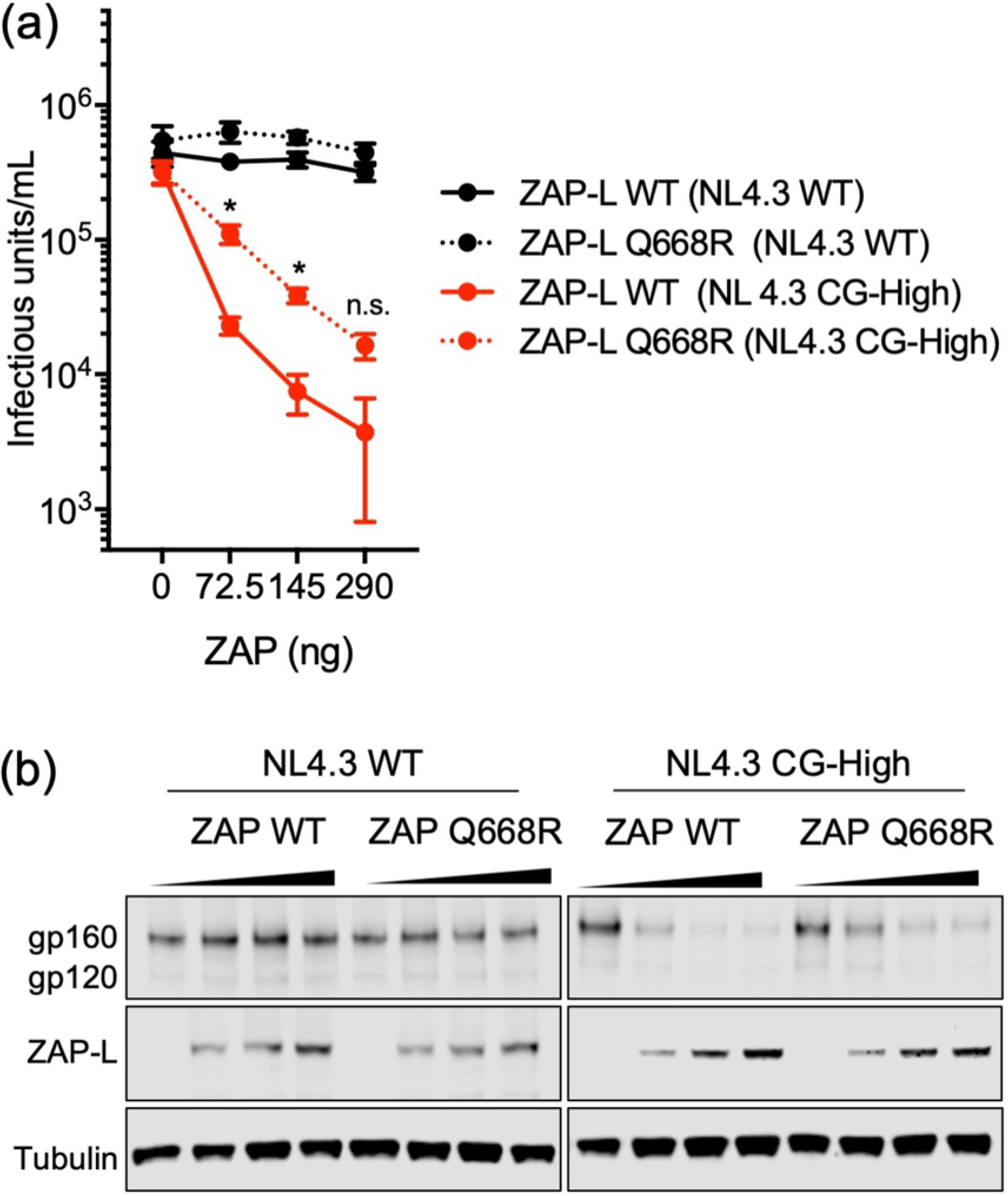
Antiviral potency of ZAP-L is reduced by the Q668R mutation. (**a**) HEK 293T *ZAP*^-/-^ *TRIM25*^-/-^ cells were transfected with a provirus of either HIV-1 (NL4.3 WT) control or a CpG-enriched mutant (NL4.3 CG-High), together with a plasmid encoding TRIM25 and increasing concentrations of a plasmid encoding WT ZAP-L or the ZAP-L Q668R mutant. After 48 hours, produced virus was harvested, filtered and titered. *, indicates p<0.05; ns, not significant. (**b**) Immunoblots of whole cell lysates showing expression levels of HIV-1 proteins (gp160 and gp120) and ZAP-L. Tubulin was used as loading control. Results are representative of two independent experiments.

## Discussion

While it is established that the N-terminal RBD of ZAP containing its first four zinc fingers is both necessary and sufficient to recognize CpG-rich viral RNA and direct their degradation, how the downstream domains contribute to ZAP’s antiviral function remains to be elucidated. Our structural and biochemical studies here reveal that the central regions of ZAP, comprising the fifth zinc finger and two WWE modules, integrate to form a single folded domain. This ZAP central domain displays an electropositive surface and features a prominent pocket in the second WWE module. Our data indicate that this pocket binds to the repeating *iso* (ADP-ribose) unit found in PAR. This work therefore provides further support for the idea that WWE domains have the general function of acting as PAR-binding modules [9, 21].

How might PAR binding relate to ZAP’s antiviral function? Being an extended polynucleotide, PAR chains have the requisite size and architecture to function as a polyvalent scaffold that facilitates clustering of binding partners [11, 26]. Multiple CpGs are required to target an mRNA strand for degradation and each ZAP RBD can only bind a single CpG dinucleotide, implying that selective recognition requires formation of a multivalent ZAP/mRNA complex [3–5]. Thus, a simple model is that PAR binding by ZAP can facilitate recognition and subsequent RNA processing by promoting local clustering of the protein molecules and thereby shifting the interaction equilibria to favor association. It is possible that such an affinity amplification mechanism may become more acutely important in certain contexts, for example when pathway components are limiting or when viral RNA levels are low.

Alternatively, PAR binding may be a means to regulate ZAP’s sub-cellular distribution. PAR is critical for the formation and maintenance of RNA stress granules [16]. ZAP is diffusely cytoplasmic, but upon viral infection is localized to stress granules [15]. ZAP was also identified as a component of granules induced by oxidative stress [27]. Similarly, at least one ZAP co-factor, TRIM25, has been reported to be associated with stress granules [27–29]. Although it remains to be established whether stress granules are the actual site of antiviral activity, it was recently reported that differential access of the long and short ZAP isoforms to target mRNA populations is regulated by sub-cellular localization [30]. Specifically, ZAP-L is targeted to intracellular compartments by a C-terminal posttranslational modification (prenylation [31]) where it can access viral mRNA, whereas ZAP-S lacks this targeting signal and remains cytosolic where it accesses a different pool of cellular mRNA [30]. Both forms of ZAP contain the central domain, and thus, binding of this domain to PAR may be an additional mechanism to regulate where and when ZAP engages its targets. It is important to note, however, that the RBD (or ZAP-N) can independently associate with stress granules [15], although it is not clear whether this occurs through direct PAR binding by the RBD or indirectly through bound RNA. Indeed, our data indicate that loss of the central domain’s PAR-binding activity only reduces – and does not eliminate – ZAP-L’s ability to inhibit virus replication. Thus, the PAR-binding function of the central domain is an ancillary activity that contributes to the overall efficiency of viral RNA recognition and degradation.

## Materials and Methods

### Plasmids

ZAP-L was obtained from Addgene (plasmid #45907). ZAP-CD was generated by using primers containing a Kozak sequence and an N-terminal HA tag, and inserted into pCDNA3-MCS between the EcoRI and NotI sites. ZAP mutants (W611A, Q668A, and Q668R) were generated by using the QuikChange Lightning site-directed mutagenesis kit (Agilent Technologies). *E. coli* expression plasmids were generated by sub-cloning from the Addgene plasmid using Gibson assembly. All coding sequences were confirmed by DNA sequencing.

### ZAP-CD purification, crystallization and structure determination

ZAP-CD was expressed with a His_6_-SUMO leader sequence in *E. coli* BL21(DE3) cells by using the autoinduction method [32]. The His-tagged fusion protein was purified by using Ni-NTA chromatography, the tag was removed with Ulp1 protease, and the untagged ZAP-CD protein was purified to homogeneity using anion exchange chromatography. The protein was exchanged into storage buffer (20 mM Tris, pH 8, 100 mM NaCl, 1 mM TCEP) by using preparative size exclusion or dialysis.

Crystallization was performed in sitting drops, by mixing protein and precipitant (0.8-1.1 M sodium nitrate, 0.1 M sodium acetate, pH 4.8-6.0) at a volume ratio of 3:1. Crystals were cryo-protected in 20% PEG 400 and diffraction data were collected at the Advanced Photon Source beamline 22-ID. The structure was solved by single anomalous diffraction methods from a selenomethione dataset, and the data quality was sufficiently high to permit automatic model building by PHENIX software [33] directly from integrated data with only some manual rebuilding required. Structure statistics are summarized in **Table 1**. At the completion of structure refinement, weak but clear difference density was observed in the second WWE domain, suggesting that some small molecule had co-purified and co-crystallized with ZAP-CD.

Indeed, the A_260_/A_280_ ratios of purified samples were ~0.6-0.7, indicating the presence of low levels of some A_260_-absorbing component. However, it was clear that only a small fraction of the protein was bound, because the co-purifying small molecule was not detected by mass spectrometry analysis of the sample used for crystallization. It appears that this bound fraction was the one that crystallized, because the crystals were very sparse and small.

### Differential scanning fluorimetry

Thermal melting profiles were measured by using a Tycho (NanoTemper), following the manufacturer’s instructions.

### Preparation of PAR

PAR was enzymatically prepared as previously described [22].

### Size exclusion binding assay

Size exclusion was performed in 20 mM Tris, pH 8, 100 mM NaCl, 1 mM TCEP. Four A_280_ absorbance units of ZAP-CD (77 μM) was mixed with equal volume of four A_260_ absorbance units of PAR (296 μM in terms of ADP-ribose subunits), incubated for 20 min at room temperature, and then injected on a Superdex 200 30/100 column and developed at a flow rate of 0.5 mL/min at room temperature. For control injections, ZAP-CD or PAR was mixed with buffer alone.

### Negative stain electron microscopy

Samples (3 μL) were onto the carbon side of carbon-coated copper grids (Electron Microscopy Sciences) for 90 s, rinsed for 90 s by flotation on a drop of water, blotted dry, stained for 90 s by floating on a drop of 2% (*w*/*v*) solution of uranyl acetate, blotted dry, and allowed to air dry. Images were viewed on an FEI Tecnai Spirit transmission electron microscope operating at 120 kV.

### Cells and plasmid transfections

HEK 293T and HeLa cell lines were maintained in DMEM supplemented with 10% FBS (fetal bovine serum), 100 U/mL penicillin, and 100 μg/mL streptomycin. Transfections were performed by using Hilymax (Dojindo Molecular Technologies) according to the manufacturer’s instructions.

### Immunoprecipitation and immunoblotting

At 48 h after transfection, HEK 293T cells were washed twice with PBS (phosphate-buffered saline) and lysed in IP buffer (50 mM Tris, pH 7.5, 150 mM NaCl, 0.2% Triton X-100) containing 1 μM ADP-HPD (Millipore) and cOmplete Mini EDTA-free inhibitor (Roche) for 20 min (with rotation at 4 °C). The lysate was clarified by centrifugation at 4 °C for 15 min at 18,800 ×*g*. A final concentration of 10 μg/mL cytochalasin B (Sigma) and 25 μM nocodazole (Sigma) were then added to the supernatant. Samples were then mixed with either anti-HA magnetic beads (Thermoscientific) or protein G magnetic beads (Thermoscientific) coupled with anti-pADPr (Abcam) (beads were pre-blocked with 1% BSA). After overnight incubation at 4 °C, the beads were washed 4 times with IP buffer, re-suspended in SDS loading buffer, boiled for 5 min, electrophoresed on a Novex 4-20% Tris-Glycine mini-gel (Invitrogen), and then transferred onto PVDF membranes. Mouse anti-poly (ADP-ribose) polymer [10H] (1:1000; Abcam), mouse anti-HA [F-7] (1:3000; Santa Cruz), mouse anti-β-Actin [C4] (1:3000; Santa Cruz), and HRP-conjugated goat antibody to mouse (1:10,000; Azure Biosystems) were used for detection.

### Immunostaining and fluorescence microscopy

At 24 h after transfection, HeLa cells were treated with or without 250 μM sodium arsenate for 30 min. HeLa cells on coverslips were washed twice in PBS and fixed for 15 min in 4% paraformaldehyde and then permeabilized with 0.5% Triton X-100. Coverslips were washed three times in PBS, then blocked with 3% BSA in TBST (Tris-buffered saline supplemented with 0.05% Tween 20) for 30 min. Cells were incubated with primary antibodies at room temperature for 1 h, and then with Alexa-Fluor-conjugated secondary antibodies and Hoechst 33342 (ThermoFisher) at RT for 30 min. Coverslips were mounted on glass slides with ProLong Gold antifade solutions (Molecular Probes). Mouse monoclonal anti-poly (ADP-ribose) polymer [10H] (1:50; Abcam) and rabbit monoclonal anti-HA [C29F4] (1:500, Cell Signaling) were used as primary antibodies. Secondary antibodies were donkey anti-mouse IgG H&L Alexa Fluor 647 (1:500; Abcam) and donkey anti-rabbit IgG H&L Alexa Fluor 488 (1:500; Abcam).

Microscopy was performed by using an LSM 880 confocal laser scanning microscope (Zeiss) with a 63× oil immersion objective (NA 1.4) and Zen imaging software (Zeiss). Images were collected simultaneously using the 405, 488 and 633 nm excitation laser lines.

### Antiviral activity assays

HEK 293T *ZAP*^-/-^ *TRIM25*^-/-^ cells were transfected with proviral plasmids of either wild type HIV-1 or a CpG-enriched mutant (NL4-3) [5], together with a plasmid encoding TRIM25 and increasing concentrations of a plasmid encoding wild type ZAP-L or mutant ZAP-L (Q668R). Cells were incubated for 48 h at 37 °C. Produced virus was then harvested, filtered and titered on MT4-GFP cells to determine infectious units per mL. Statistical significance was assessed using multiple *t*-tests (wild type ZAP-L versus Q668R ZAP-L, wild type and CpG-enriched NL4.3 viruses, corrected for multiple comparisons using the Holm-Sidak method) and two-way ANOVA.

## Acknowledgements

We thank Jacint Sanchez for Jonathan Wagner assistance and advice with protein purification, crystallization and diffraction data collection. This work was funded by National Institutes of Health Grants R01-AI150479 (O.P.), U54-AI150470 (O.P., B.K.G.-P. and P.D.B.) and R01-AI150111 (P.D.B.).

